# Inducible Plasmid Self-Destruction (IPSD) assisted genome engineering in lactobacilli and bifidobacteria

**DOI:** 10.1101/649921

**Authors:** Fanglei Zuo, Zhu Zeng, Lennart Hammarström, Harold Marcotte

**Author notes:** Corresponding Author: E-mail: F.Z. and H.M.

## Abstract

Genome engineering is essential for application of synthetic biology in probiotics including lactobacilli and bifidobacteria. Several homologous recombination system-based mutagenesis tools have been developed for these bacteria but still, have many limitations in different species or strains. Here we developed a genome engineering method based on an inducible self-destruction plasmid delivering homologous DNA into bacteria. Excision of the replicon by induced recombinase facilitates selection of homologous recombination events. This new genome editing tool called Inducible Plasmid Self-Destruction (IPSD) was successfully used to perform gene knock-out and knock-in in lactobacilli and bifidobacteria. Due to its simplicity and universality, the IPSD strategy may provide a general approach for genetic engineering of various bacterial species.

---

Lactobacilli and bifidobacteria, have been extensively used as probiotics and have become increasingly studied as living diagnostics and therapeutics that can probe and improve human health.^1-4^ In contrast, the genetic tools (especially mutagenesis) to investigate and enhanced their probiotic activity are rather poorly developed, especially for bifidobacteria.^5^ The traditional genome engineering methods are largely dependent on the bacterial transformation efficiency. This defect can be overcome by the conditional replicate plasmid assisted homologous recombination, such as plasmids containing a thermosensitive replication origin,^6, 7^ but the latter may have a limited host range. Some counterselection systems based on *upp* and *pyrE* genes has been developed.^8, 9^ However, this strategy needs modification of the parental strains and could not resolve the problem of the low frequency of single-crossover homologous integration events. Prophage recombinase-assisted double-stranded (dsDNA) recombineering can be used for marker-free genome engineering, but it is not a seamless strategy since a “scar” is left at the modification site after excision of the selection marker.^10^ ssDNA recombineering allows high efficiency mutagenesis in lactobacilli and lactococci independent of antibiotic selection, while it still requires high transformation efficiency (i.e. ∼≥10^5^ cfu/μg DNA) and inducible expression of recombinase RecT.^11^ Therefore, it might be difficult to be used in strains with low transformation efficiency. More recently, CRISPR-Cas system has been applied for genome editing in various lactic acid bacteria species, including *L. reuteri*,^12^ *L. casei*,^13^ *L. plantarum*,^14^ and *L. lactis*.^15^ However, the editing outcomes can vary across strains and between methods,^14^ and have occasional target failure.^16^ Furthermore, CRISPR-Cas system has not yet been successfully used in bifidobacteria.^17^ Therefore, more flexible and universal genome engineering strategies need to be developed for a better understanding and application of these health-promoting microorganisms.

We have designed a novel vector that could be destructed under certain conditions facilitating the selection of homologous recombination event between the vector and chromosomal DNA. Essentially, the replicon and the antibiotic resistance gene were separated by two oriented *six* or *lox* fragments.^18, 19^ Upon addition of the inducer and expression of the site-specific recombinase (β or Cre),^18, 19^ the vector can recombine and lose function due to the excision of the replicon (Figure 1A). This vector termed Inducible Plasmid Self-Destruction (IPSD) could be used to assist bacterial genome engineering, including gene deletion, insertion, and replacement (Figure 1B). When the vector delivering homologous DNA into bacteria, homologous recombination naturally occurred, induce plasmid destruction could facilitate homologous recombination and selective enrichment of single-crossover integrant precursor (Figure 1B).

**Figure 1.**
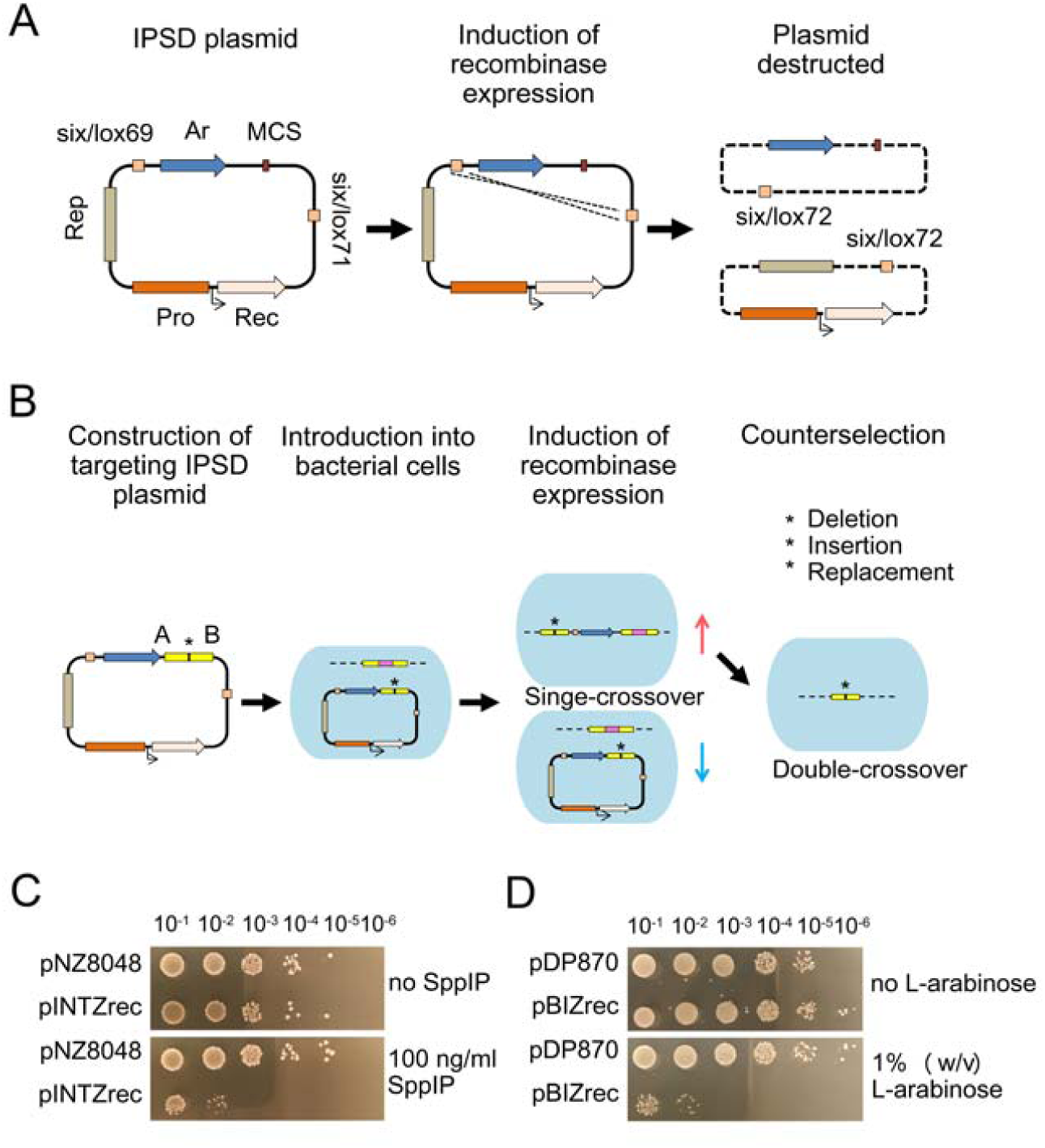
IPSD strategy for bacterial genome engineering. (A) Schematic illustration of inducible plasmid self-destruction. A vector in which the replicon and the antibiotic resistance gene are separated by two oriented *six* or *lox* fragments was constructed. Upon addition of the inducer and β or Cre mediated recombination, the vector loses function due to the excision of the replicon. Rep, replicon; Pro, controlled expression promoter; Rec, recombinase; six/lox: two oriented *six* or *lox* target sequence sites; Ar, antibiotic resistance gene; MCS, multiple cloning sites. (B) Schematic illustration of IPSD assisted bacterial genome engineering, including gene deletion, insertion, and replacement (indicated by an asterisk). After recombination, the ratio of subpopulation harboring the episomal vector decreased (blue arrows) while the ratio of subpopulation with integrated DNA fragment through single-crossover increased (red arrows) under antibiotic selection. The single-crossover integrant precursor could be screened by PCR and the double-crossover clones could be selected by counterselection. The homologous ends flanking the target gene are indicated by A and B. The target gene on the chromosome is represented by a pink rectangle. (C) Growth of the indicated *L. gasseri* DSM 14869 strain on MRS agar plate supplemented with 10 μg/ml chloramphenicol and in presence or not of 100 ng/ml SppIP. (D) Growth of the indicated *B. longum* NCC 2705 strain on MRSc agar plate supplemented with 100 μg/ml spectinomycin and in presence or not of 1 % (w/v) L-arabinose.

As a proof-of-concept, an IPSD plasmid pINTZrec was constructed from pNZ8048 based on the β-*six* site specific recombination system (Figure S1a) in which the *β* recombinase gene was under the control of the sakacin-inducible promoter P_orfX_. When the plasmid pINTZrec carrying a chloramphenicol resistance gene was introduced into the vaginal probiotic strain *L. gasseri* DSM 14869,^20^ the pINTZrec transformant showed a dramatic sensitivity to Sakacin P (SppIP) induction. The viability of SppIP-induced pINTZrec transformant on MRS agar plates supplemented with chloramphenicol was decreased by three to four orders of magnitude compared to the non-induced control (Figure 1C), suggesting that the plasmid pINTZrec was destructed following expression of *β* recombinase. Such observation was also found in other *Lactobacillus* species, such as *L. paracasei, L. acidophilus*, and *L. plantarum* (Figure S2). We further extended this method to *Bifidobacterium*, a genus for which genetic tools are poorly developed. An IPSD plasmid pBIZrec was constructed from pDP870 based on the Cre-*lox* site specific recombination system (Figure S1b) and in which the recombinase gene *Cre* was under the control of the L-arabinose-inducible promoter araC-P_BAD_. The plasmid pBIZrec carrying a spectinomycin resistance gene was introduced into *B. longum* NCC 2705 and the pBIZrec transformant also showed decreased viability on agar plates supplemented with spectinomycin following cultivation in the presence of L-arabinose, by four to five orders of magnitude compared to the non-induced control (Figure 1D). These results indicated that the IPSD strategy could be used for assisting genome engineering in lactobacilli and bifidobacteria.

Our previous work showed that the transformation efficiency of *L. gasseri* DSM 14869 was very low possibly due to the thickness of EPS covering the cell surface or the presence of two resident plasmids, and therefore, generating mutation in this strain using existing methods was not possible ^20^ (unpublished data). In order to demonstrate IPSD assisted genome engineering in *L. gasseri* DSM 14869, we targeted *upp*, a non-essential gene encoding uracil phosphoribosyltransferase (UPRTases) that is commonly used as a counterselection marker.^8^ Arecombinant plasmid pINTZrec-Δupp containing homologous regions up-and downstream of *upp* gene was constructed and introduced into *L. gasseri* DSM 14869. The transformant was induced by SppIP and the single-crossover integration event was selected using colony PCR (Figure 2A and Figure S3a). Ten out of randomly picked 28 (36%) colonies showed absence of plasmid pINTZrec-Δupp, suggesting that the Cm^r^ expression cassette was integrated into the chromosome of *L. gasseri* DSM 14869 by a single-crossover event. These results were further confirmed by PCR on DNA extracted from isolated clones and six clones showed correct integration (Figure S3b). Following growth of the single-crossover clones in the absence of antibiotics, the double-crossover *upp* mutant could easily be selected by counterselection of Cm^s^ colonies or by selection of colonies resistant to 5-Fluorouracil (5-FU) (Figure 2B and Figure S3c). The *L. gasseri* DSM 14869 *upp* mutant showed resistance to 5-FU (100 µg/ml) in contrast to the parent strain due to the abolished conversion of 5-FU into cell toxic 5-fluorodeoxyuridine monophosphate (5-FdUMP) (Figure 2C).^8^ We have also generated several mutants of cell surface property related genes and integrative expression of broad and potent HIV-1-neutralizing antibodies in *L. gasseri* DSM 14869 based on this method (unpublished data). These results suggest that the IPSD plasmid could efficiently be used for genome engineering in lactobacilli.

**Figure 2.**
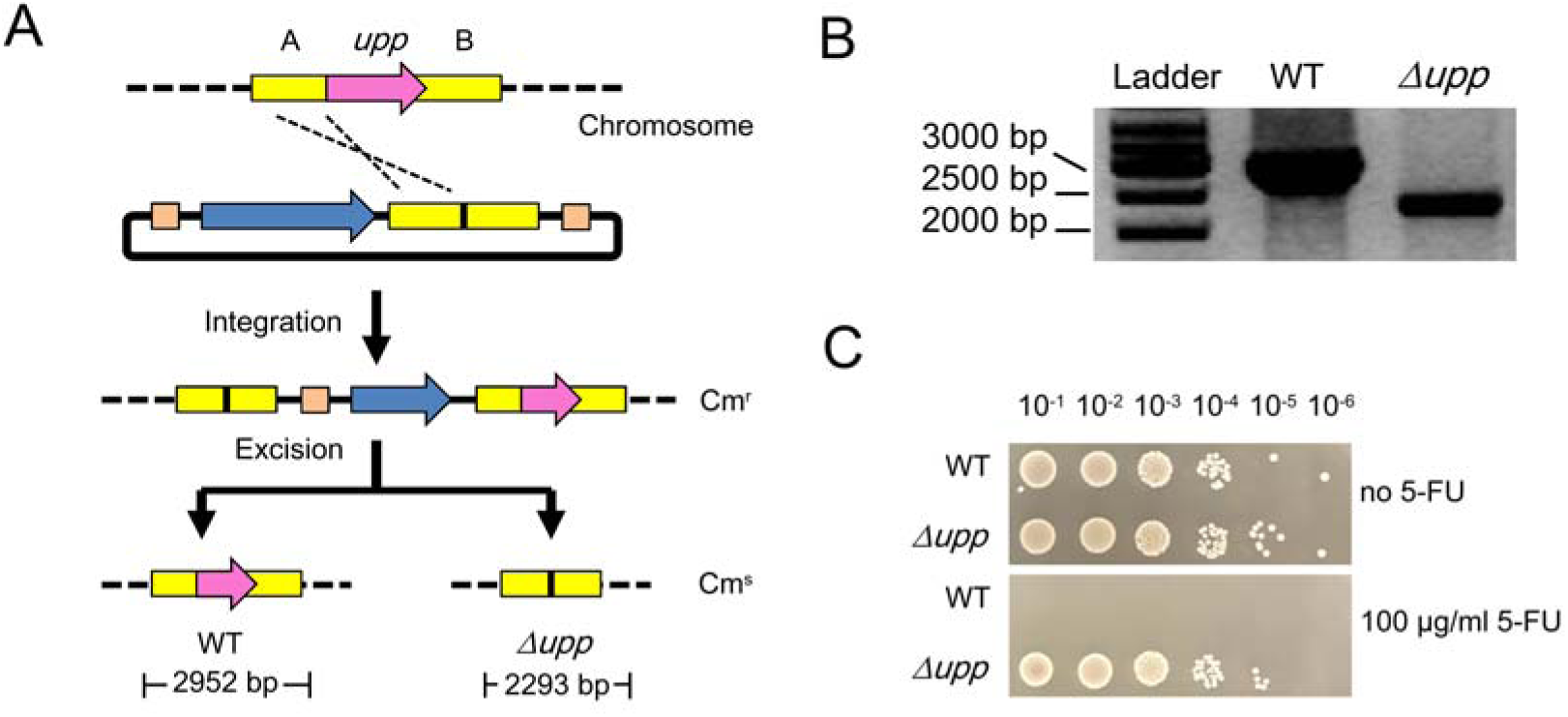
Generation of *L. gasseri* DSM 14869 *upp* deletion mutant. (A) Schematic representation of a two-steps homologous recombination to obtain a *upp* deletion mutant. (B) The *L. gasseri* DSM 14869 *upp* gene was deleted and the deletion mutant verified by PCR using primer pairs uppleft-F and uppright-R. (C) The *L. gasseri* DSM 14869 *upp* mutant showed resistance to 100 μg/ml of 5-FU compared to the WT. Overnight liquid cultures of the *L. gasseri* DSM 14869 WT and *upp* mutant were adjusted at the same OD_600_ before serial 10-fold dilutions were spotted on plates with or without 5-FU. The plates were incubated at 37 °C and photographed the next day.

We subsequently tested this method in *Bifidobacterium*. The *tetW* gene provides tetracycline resistance to dominant bifidobacterial species from the human gastrointestinal tracts, and it might be horizontally transferred to other species.^21^ Concerning the safety of probiotics, gene deletion could be the best way to prevent the potential horizontal transfer of functional antibiotic resistant genes. The *tetW* gene from different bifidobacterial strains shared high identity but with large variation in their flanking regions.^22^ Therefore, we inactivated the *tetW* gene by in-frame deletion in *B. longum* IF3-53, an infant feces isolate showing tetracycline resistance ^23^ (Figure 3A). An IPSD plasmid pBIZrec-ΔtetW was constructed and transformed into *B. longum* IF3-53. The transformant was induced by L-arabinose and the single-crossover integration event was selected using colony PCR (Figure 3B and Figure S4a). Three out of 16 (19%) randomly picked colonies were shown to contain the correct insertion (Figure S4b). The double-crossover *tetW* mutant was subsequently selected by counterselection of Sm^s^ colonies (Figure 3B). The mutant was sensitive to tetracycline compared to the parent strain, as the minimum inhibitory concentration (MIC) was reduced from 32 µg/ml to 1 µg/ml (Figure 3C). The results suggest that the IPSD plasmid could be used for genome engineering in bifidobacteria as well.

**Figure 3.**
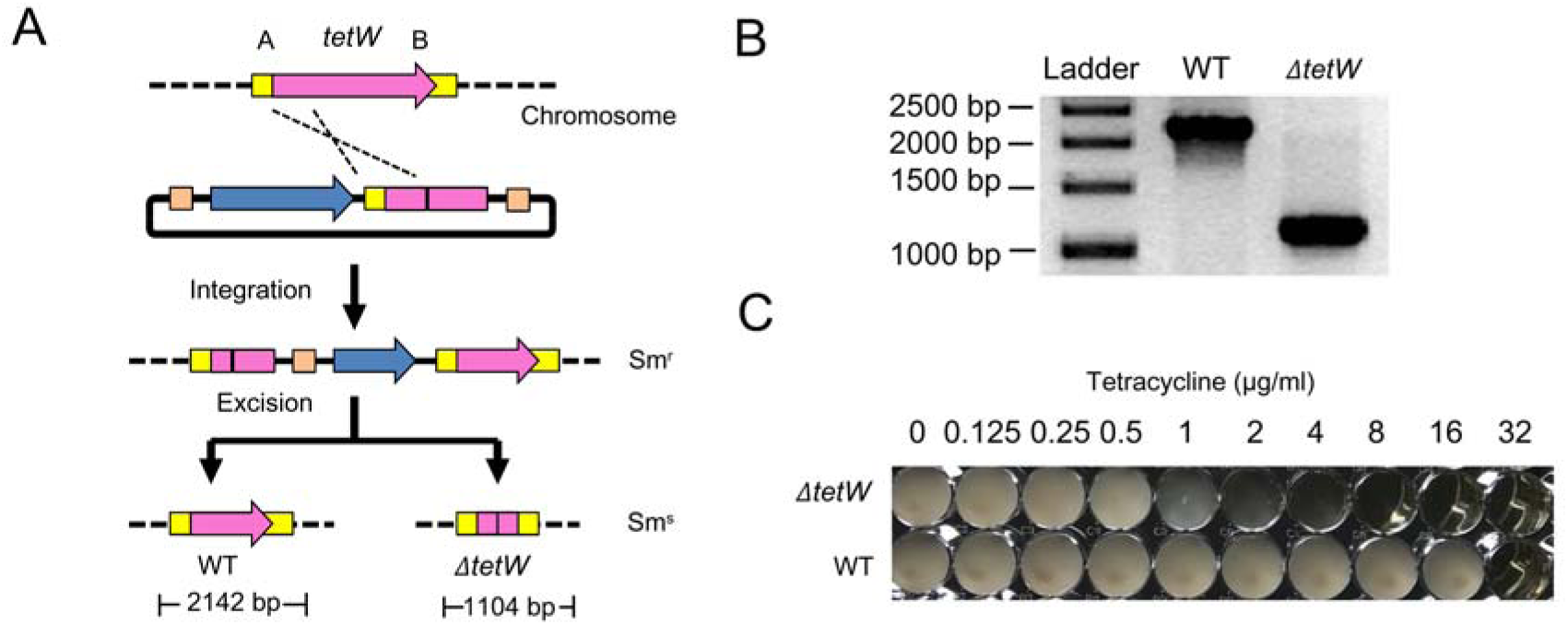
Generation of *B. longum* IF3-53 *tetW* in-frame deletion mutant. (A) Schematic representation of a two-steps homologous recombination to obtain a *tetW* in-frame deletion mutant. (B) The *B. longum* IF3-53 *tetW* gene was disrupted and the mutant verified by PCR using primer pairs tetWleft-F and tetWright-R. (C) The minimum inhibitory concentration (MIC) of tetracycline was lower for *B. longum* IF3-53 *tetW* mutant compared to WT. Overnight liquid cultures of the *B. longum* IF3-53 WT and *tetW* mutant were adjusted at the same OD_600_ before added into 96-well plates containing two-fold serial dilutions of tetracycline hydrochloride. The plates were incubated at 37 °C under anaerobic conditions and photographed the next day.

We further tested the IPSD vector for stable chromosomal integration and expression of heterologous genes in *Bifidobacterium*. A gene *LpKatL* encoding catalase from *L. plantarum* was inserted into the chromosome of *B. longum* NCC 2705 under the control of the native P_hup_ promoter (Figure 4A and 4B). This integration site was selected based on several criteria as reported previously.^24^ First, *LpKatL* gene was integrated downstream of gene *hup*, encoding the histone-like protein HU, which has high and constitutive expression level.^25^ Polycistronic expression of *LpKatL* was driven by the promoter of the upstream gene *hup*. Second, an optimized ribosome-binding site (RBS) (AAGGAG) and a 5 nt spacer length was introduced in front of *LpKatL* gene for efficient translation. Finally, the insertion location between a stop codon and a terminator was selected to minimize the polar effects. The recombinant strain showed catalase activity in the presence of hematin (Figure 4C), and improved viability when exposed to H_2_O_2_ as compared to the parent strain (Figure 4D).

**Figure 4.**
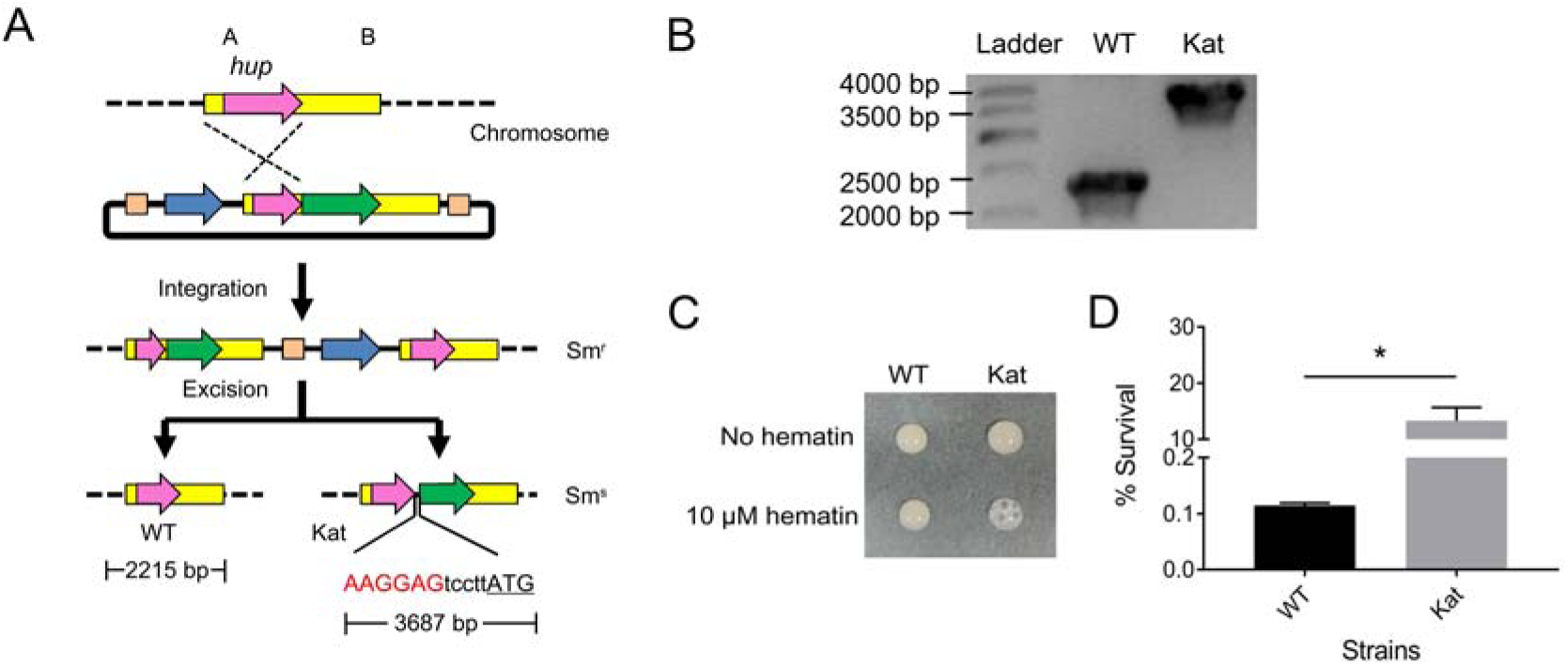
Integrative expression of *Kat* gene in *B. longum* NCC2705. (A) Schematic representation of a two-steps homologous recombination to obtain a integrative *LpKatL* expressing mutant. The catalase gene *LpKatL* was inserted into the chromosome of *B. longum* NCC 2705 and placed under the control of P_*hup*_ by a polycistronic structure, an optimized RBS was introduced (marked in red), the start codon is underlined. (B) The *LpKatL* integrative mutant verified by PCR using the primer pair Katleft-F and Katright-R. (C) Catalase activity test. O_2_ bubble was formed upon addition of H_2_O_2_ on the cell pellet of the LpKatL recombinant *B. longum* NCC 2705 strain harvested from liquid culture in MRS broth supplemented with hematin. (D) The viability of the LpKatL recombinant *B. longum* NCC 2705 strain was dramatically improved compared to the WT under H_2_O_2_ challenge. Values are averages from two independent experiments, ±standard deviation. *p*<0.05.

The major advantage of this method is that it does not depend on transformation (or conjugation) efficiency. However, it needs two pre-requirements: 1) a functional replicon allowing the plasmid to replicate in the host bacteria; 2) a tightly controlled expression element used to drive recombinase gene expression. The plasmid pINTZrec contains a pSH71 replicon which is broad-host-range and high-copy-number widely used for lactobacilli plasmid construction.^26^ The plasmid pBIZrec contains a pNCC293 replicon which could replicate in at least seven different bifidobacterial species.^27, 28^ So these two IPSD plasmids are expected to be universally used for genome engineering in different lactobacilli and bifidobacteria species. However, the inducible promoters used in this study are not effective in some strains, either due to strong background expression (*L. sakei* NC03) (Figure S5a) or low induced expression (*L. rhamnosus* GG, *B. lactis* Bb12) (Figure S5b and S5c). Therefore, the development of a universal tightly controlled expression element for lactobacilli or bifidobacteria, such as tetracycline-regulated systems, is necessary.^29^ Another disadvantage is that the IPSD strategy only addressed the selection of single-crossover integration while the double-crossover excision of the antibiotic resistant gene was achieved by traditional counterselection of antibiotic sensitive clones. Combination with various counterselection methods, especially CRISPR-Cas-assisted negative selection, will further improve the genome engineering efficiency in bacteria.^12^

In conclusion, we have shown that the IPSD plasmid can be used for genome engineering in lactobacilli and bifidobacteria, with the potential to be extended to other bacterial species. The IPSD strategy could be used in a range of synthetic biology applications in both the food and pharmaceutical industry such as identification of probiotic genes, metabolic engineering, and delivery of therapeutics,^30-32^ thus opening new avenues for the engineering of biotherapeutic agents with enhanced health-promoting functional features.

## METHODS

### Bacterial strains, plasmids and growth conditions

Bacterial strains and plasmids used in this study are listed in Table S1. *Lactobacillus* strains were generally cultured at 37 °C in deMan Rogosa Sharpe (MRS) medium (Difco, BD BioSciences). For *upp*-based double-crossover event selection, the *Lactobacillus* strain was grown on semi-defined medium (SDM) agar plates.^33^ *Bifidobacterium* strains were cultured in MRS supplemented with 0.05 % (w/v) L-cysteine HCl (MRSc) at 37 °C under anaerobic conditions. *Escherichia coli* strains were cultured in Luria-Bertani broth at 37 °C with rotary shaking at 200 rpm, or on LB agar plates. When needed, antibiotics were supplemented at the following concentrations: 10 μg/ml chloramphenicol for *Lactobacillus* strains and *E. coli* VE7108 strain, and 100 μg/ml spectinomycin for *Bifidobacterium* strains and *E. coli* DH5α strain.

### Plasmid construction

The primers used in this study are listed in Table S2. For construction of IPSD vector pINTZrec used for lactobacilli genome engineering, the two *six* DNA fragments were amplified from site-specific integration vector pEM76 by using the primers pairs SIX-F1&R1 and SIX-F2&R2, respectively, and inserted into plasmid pNZ8048 on each flank of the Cm^r^ expression cassette generating pNZ8048-SIX. The orientation of the insertion was confirmed by sequencing. Multiple cloning sites was introduced into pNZ8048-SIX by inserting a linker between *Pst*I and *Bgl*II, resulting in the plasmid pNZmcs-SIX. The *β*-recombinase gene was amplified from plasmid pEM94 and inserted into plasmid pVPL3017 downstream of the sakacin-inducible promoter P_orfX_ generating pVPL3017-rec. Subsequently, the P_orfX_-rec expression cassette was digested using *Sal*I and *Hind*III, and inserted into similarly digested pNZmcs-SIX plasmid to obtain the final plasmid pINTZrec.

For construction of the plasmid used for *L. gasseri* DSM 14869 *upp* gene deletion, two 1090 bp DNA fragments, upstream and downstream of *upp* gene, were amplified from genomic DNA of *L. gasseri* DSM 14869 using the primer pairs upp-up-F/upp-up-R and upp-down-F/upp-down-R, respectively. The upstream DNA fragment was inserted into pMD19-T Simple vector by TA cloning generating pMD19-upp-up, then the downstream DNA fragment was inserted into pMD19-upp-up between *Sac*I and *Sph*I, generating pMD19-Δupp.Δupp was digested with *Apa*I and *Sph*I and inserted into similarly digested pINTZrec generating pINTZrec-Δupp.

For construction of IPSD vector pBIZrec used for bifidobacteria genome engineering, the *lox66-Smr-lox71* DNA fragment was amplified by PCR amplification from pDP870 using primers lox66-F and lox71-R, followed by insertion into pDP870 between *EcoR*V and *EcoR*I generating pDP870-lox. The L-arabinose inducible promoter araC-P_BAD_ was amplified from *E. coli* DH5α genomic DNA using primers araP-Cre-F1 and araP-Cre-R1 and the recombinase gene *Cre* was amplified from plasmid pAdTrack-Cre using primers araP-Cre-F2 and araP-Cre-R2. The *araP-Cre* DNA fragment was generated by overlap PCR using primers araP-Cre-F1 and araP-Cre-R2, and a mixture of *araC-P*_*BAD*_ and *Cre* PCR products as templates. The *araP-Cre* DNA fragment was then inserted into pDP870-lox between *Acc*I and *Hind*III generating pBIZrec.

For construction of the plasmid used for *B. longum* IF3-53 *tetW* gene inactivation, two 497 bp DNA fragments, upstream and downstream of *tetW* gene were amplified from *B. longum* IF3-53 genomic DNA using the primer pairs tetW-up-F/tetW-up-R and tetW-down-F/tetW-down-R, respectively. These two DNA fragments were fused by overlap PCR using primers tetW-up-F and tetW-down-R, generating ΔtetW and inserted into pBIZrec between *Apa*I and *Nhe*I resulting in pBIZrec-ΔtetW.

For construction of the plasmid used for insertion of the *LpKatL* gene into the chromosome of *B. longum* NCC 2705, the P_*hup*_-Kat DNA fragment was generated by overlap PCR. Briefly, two ∼1080 bp DNA fragments, upstream and downstream of *B. longum* NCC 2705 *hup* gene stop codon were amplified using primer pairs hup-up-F and hup-up-R, and hup-down-F and hup-down-R, respectively. *LpKatL* gene was amplified from plasmid pDP401-LpKatL using primers Kat-F and Kat-R, also introducing an optimized ribosome binding site (RBS) (AAGGAG) and a 5 nt spacer length in front of *LpKatL* gene.^34^ These three DNA fragments were fused by overlap PCR using primers hup-up-F and hup-down-R, generating P_*hup*_-Kat and inserted into pBIZrec between *Apa*I and *Nhe*I resulting in pBIZrec-P_hup_-Kat.

### Transformation

The plasmids pINTZrec and pINTZrec-Δupp were electrotransformed into *L. gasseri* DSM 14869 and other *Lactobacillus* strains according to De Keersmaecker et al.^35^ The plasmids pBIZrec, pBIZrec-ΔtetW and pBIZrec-P_hup_-Kat were electrotransformed into *B. longum* NCC 2705 and other *Bifidobacterium* strains as described previously.^27^ The plasmids pNZ8048 and pDP870 were used as controls. The transformants were confirmed by colony PCR followed by PCR on DNA extracted from pure cultures.

### Recombineering

The single colonies of *L. gasseri* DSM 14869 harboring the plasmid pINTZrec or its derivatives were grown overnight in MRS broth containing 10 μg/ml chloramphenicol. The cultures were inoculated (1 %, v/v) into MRS broth without antibiotics and grown at 37 °C until OD_600nm_ reached ∼0.30, then supplemented with 100 ng/ml sakacin P (SppIP) (Genscript). The cultures were allowed to grow overnight, and serial dilutions were plated on MRS agar supplemented with 10 μg/ml chloramphenicol and 100 ng/ml SppIP. The single-crossover events were detected by colony PCR followed by PCR on DNA extracted from pure cultures. For double-crossover selection, the single-crossover clones were grown overnight in MRS broth in absence of antibiotics, followed by spreading serial dilutions on MRS agar or SDM agar supplement of 100 μg/ml 5-Fluorouracil (5-FU) (Sigma). The colonies from MRS agar were replicated to MRS agar containing 10 μg/ml chloramphenicol, the Cm^s^ colonies were selected and detected by PCR on extracted DNA.

The single colonies of *B. longum* NCC 2705 harboring plasmid pBIZrec or its derivatives were grown overnight in MRSc broth containing 100 μg/ml spectinomycin. This overnight culture was used to inoculate MRSc broth supplemented with 1 % (w/v) L-arabinose (Sigma) without antibiotic followed by overnight culture. The cultures were serially diluted and plated on MRSc agar supplemented with 100 μg/ml spectinomycin and 1 % L-arabinose. The single-crossover events were detected by colony PCR followed by PCR on extracted DNA. For double-crossover selection, the single-crossover clones were grown overnight in the MRSc broth in the absence of antibiotics, followed by spreading serial dilutions on MRSc agar. The colonies from the MRSc agar were replicated to MRSc agar containing 100 μg/ml spectinomycin. Subsequently, the Sm^s^ colonies were selected and the double-crossover events were identified using PCR on extracted DNA.

All the mutants generated in this study were confirmed by PCR and sequencing.

### MIC assay

Minimal inhibitory concentration (MIC) was determined by the broth microdilution protocol.^36^ Overnight cultures of *B. longum* IF3-53 and *ΔtetW* mutant were harvested and re-suspended to an OD_600nm_ = 0.8 and further diluted 100-fold in MRSc. Working solutions of tetracycline hydrochloride (Sigma) were prepared in MRSc and 100 μl of two-fold dilution series were distributed in 96-well plates. Subsequently, 100 μl of bacterial suspensions were transferred to the wells. The plates were incubated under anaerobic conditions at 37 °C for 20 h. The MIC was defined as the lowest concentration of the antimicrobial agent that inhibits visible growth of the tested strains as observed with the naked eye.

### Catalase activity and H_2_O_2_ stress assay

*B. longum* NCC 2705 and recombinant *B. longum* NCC 2705 P_*hup*_::*LpKatL* were grown in MRS broth supplemented with 10 μM hematin (Sigma) for 8 h. Bacteria cells were then resuspended in TES buffer (50 mM Tris-Cl, 30 mM EDTA, 25% (w/v) sucrose, pH 8.0). Catalase activity was examined by detecting bubble formation upon addition of 10 % H_2_O_2_ to the cell suspension.^37^

For H_2_O_2_ stress, an aliquot of bacterial cultures were treated with 5 mM H_2_O_2_ for 1 h at 37 °C. Then, H_2_O_2_ was removed by washing pellets two times with PBS, and viable cells were counted by plating appropriate dilutions on MRS agar after two days anaerobic incubation at 37 °C. Cultures not treated with H_2_O_2_ were used as a reference to calculate the survival rate. Statistical analysis was performed by using two-tailed unpaired t-tests. *p*<0.05 was considered to be statistically significant.

## ASSOCIATED CONTENT

### Supporting Information

Supplementary Figures S1-S5; Supplementary Tables S1-S2.

## AUTHOR INFORMATION

### Author Contributions

F.Z. conceived the study. F.Z. and H.M. designed the experiments. F.Z. performed all the experiments with assistance from Z.Z.. F.Z., H.M. and L.H. wrote the manuscript, with contributions from all other authors. All of the authors read and approved the final manuscript.

### Notes

The authors declare the following competing financial interest: a provisional patent application covering some parts of the information contained in this article has been filed.

## ACKNOWLEDGEMENT

*E. coli* VE7108 strain was a gift from D. Mora (University of Milan, Milan, Italy). Plasmid pDP870 and *B. longum* NCC 2705 were a gift from S. Duboux (Nestec Ltd., Nestlé Research Center Lausanne, Lausanne, Switzerland). Plasmid pAdTrack-Cre was a gift from Z. Zukowska (Georgetown University Medical Center, Washington, DC, USA). This work was supported by the Swedish Research Council (Vetenskapsrådet) [U-Forsk grant 348-2013-6609 to L.H.]; and the Stiftelsen Läkare mot AIDS Forskningsfond [Fob2016-0008 to F.Z.].

