## Supporting information for "Inducible Plasmid Self-Destruction (IPSD) assisted genome engineering in lactobacilli and bifidobacteria"

^3^Present address: Department of Molecular Biosciences, The Wenner-Gren Institute, Stockholm University, Stockholm SE-10691, Sweden.

^4^Present address: College of Biotechnology, Southwest University, Chongqing 400715, P. R. China.

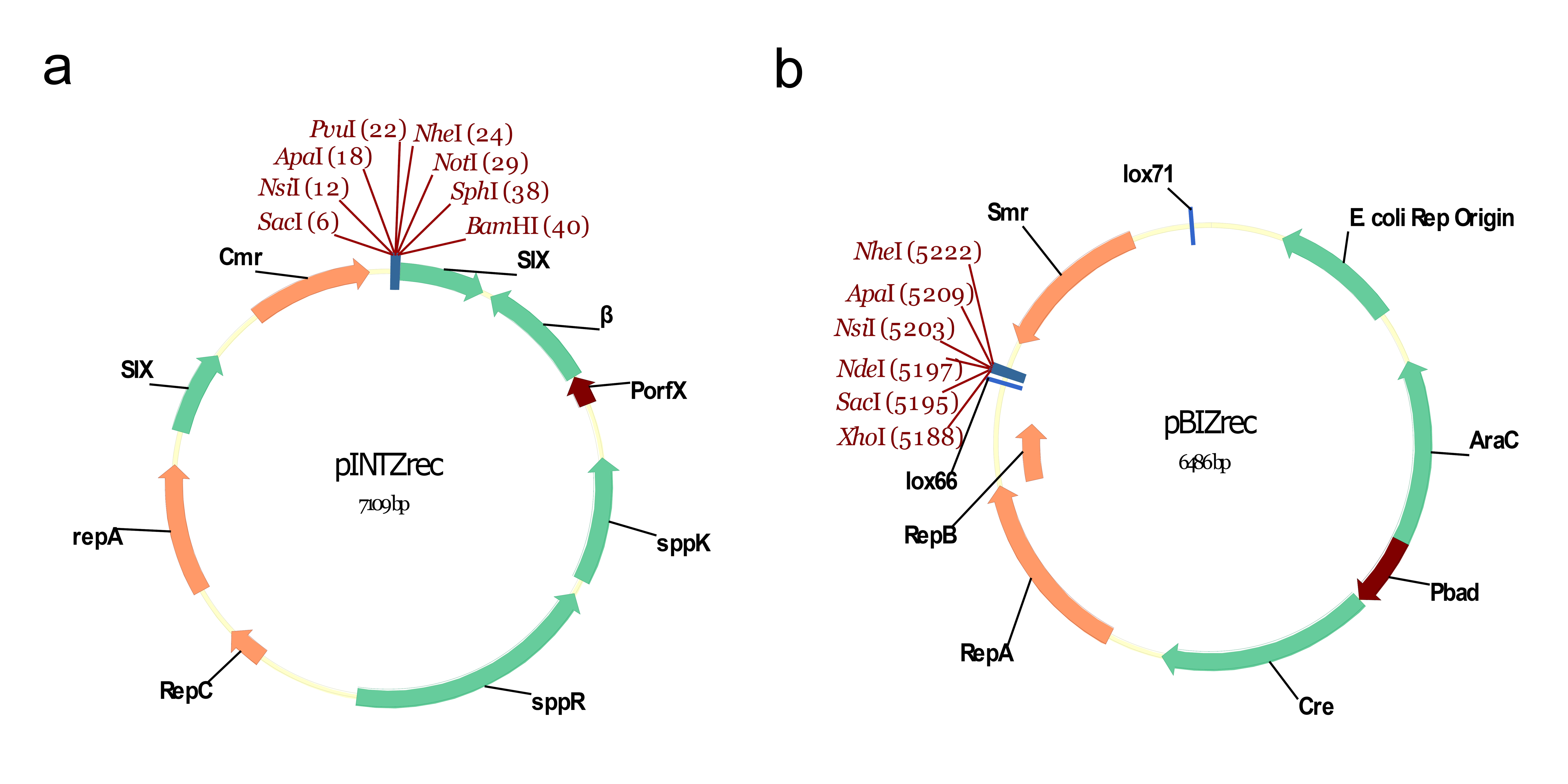

Supplementary Figure 1. The physical map of IPSD vector pINTZrec (a) and pBIZrec (b). Cmr, chloramphenicol resistance gene; Spec, spectinomycin resistance gene; Rec, recombinase gene *β*; Cre, recombinase gene *Cre*; Rep Origin, *E. coli* pBR322 ori.

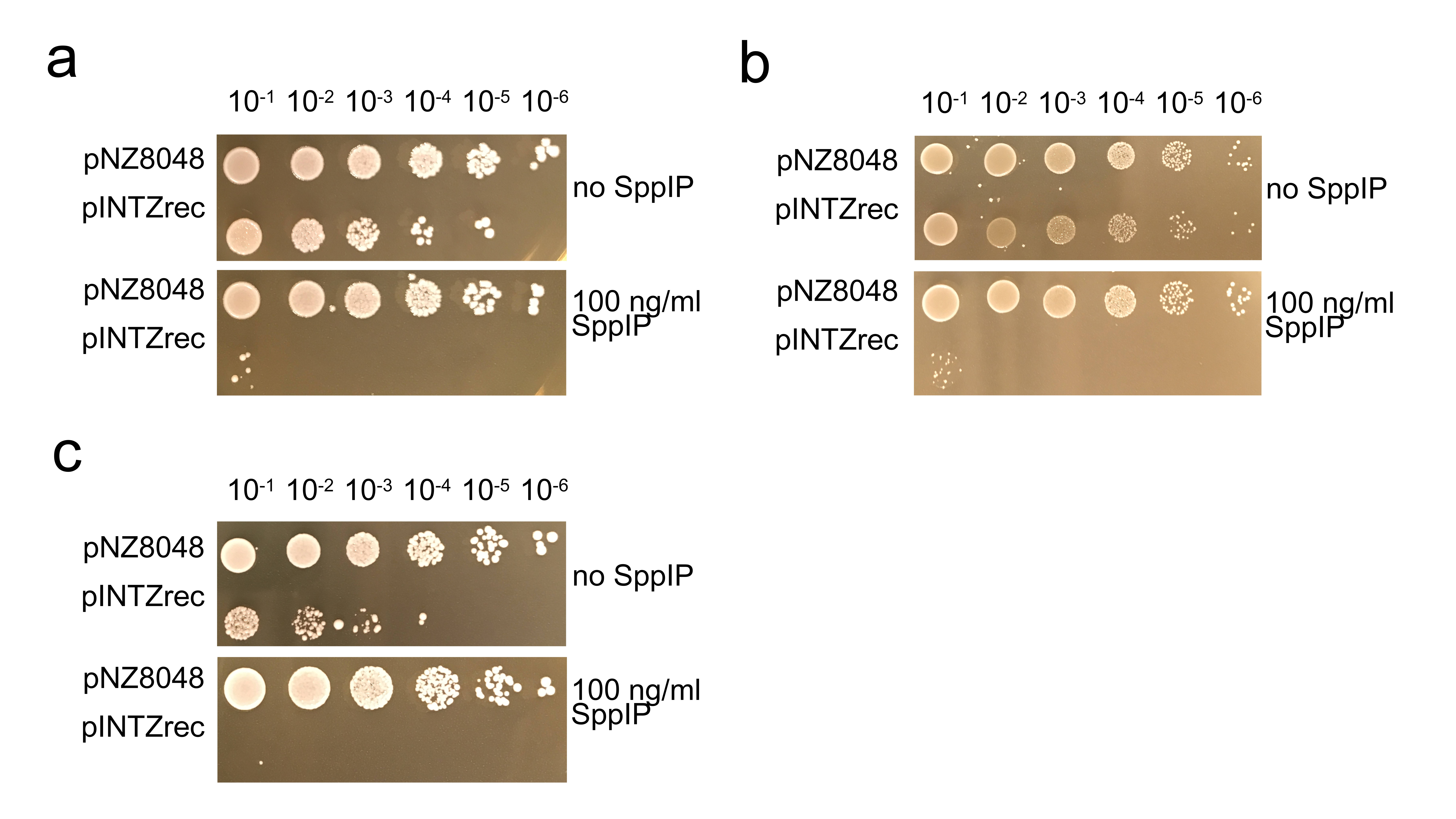

Supplementary Figure 2. Growth of the indicated *Lactobacillus* strains on MRS agar plate supplemented with 10 μg/ml chloramphenicol and in presence or not of 100 ng/ml SppIP. (a) *L. paracasei* BL23, (b) *L. acidophilus* ATCC 4356, (c) *L. plantarum* NL42. Overnight liquid cultures were adjusted at the same OD_600_ before serial 10-fold dilutions were spotted on plates with indicated supplements. The plates were incubated at 37 °C and photographed the next day.

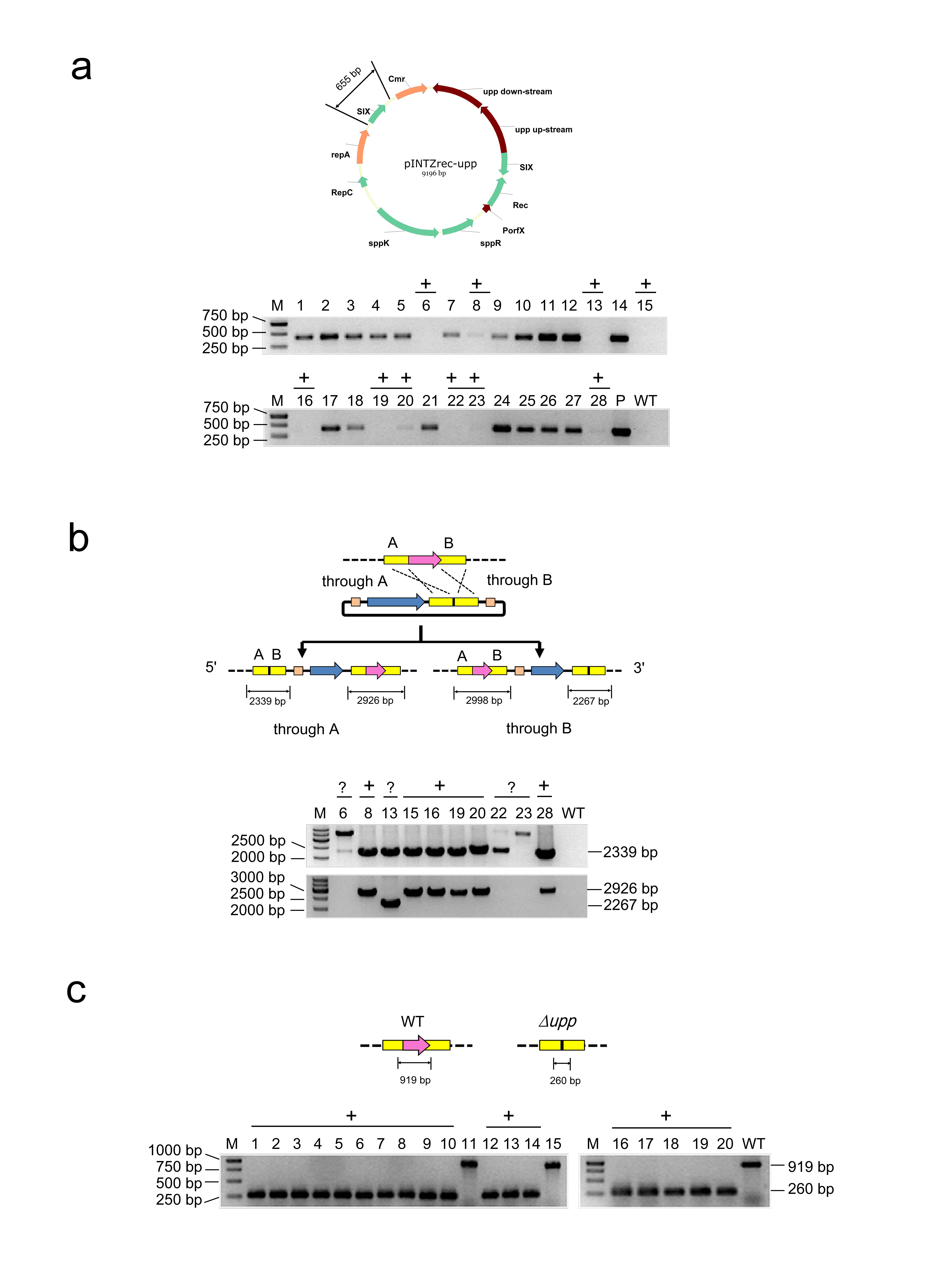

Supplementary Figure 3. Deletion of the *upp* gene in *L. gasseri* DSM 14869 by IPSD assisted genome engineering. (a) Single-crossover integration of the *upp-Cmr* fragment into the chromosome of *L. gasseri* DSM 14869. Colony PCR using primers pIrec-F and pIrec-R was performed to detect the intact plasmid pINTZrec-upp. Ten colonies without the corresponding band were regarded as candidate single-crossover clones, which were labelled "+". (b) The single-crossover event in the ten candidate clones was further confirmed by DNA extraction and PCR using primers uppleft-F and pIrecSC-R (the upper gel), or pIrecSC-F and uppright-R (the lower gel). The correct clones were labelled "+" and the uncertain clones were labelled "?". (c) Colony PCR using primers uppseq-F and uppseq-R was performed to detect *upp* double-crossover deletion mutants selected on SDM medium supplemented with 5-FU. M, GeneRuler^TM^ 1kb DNA ladder; P, DNA of *L. gasseri* DSM 14869 harboring plasmid pINTZrec-upp; WT, DNA of wild type *L. gasseri* DSM 14869.

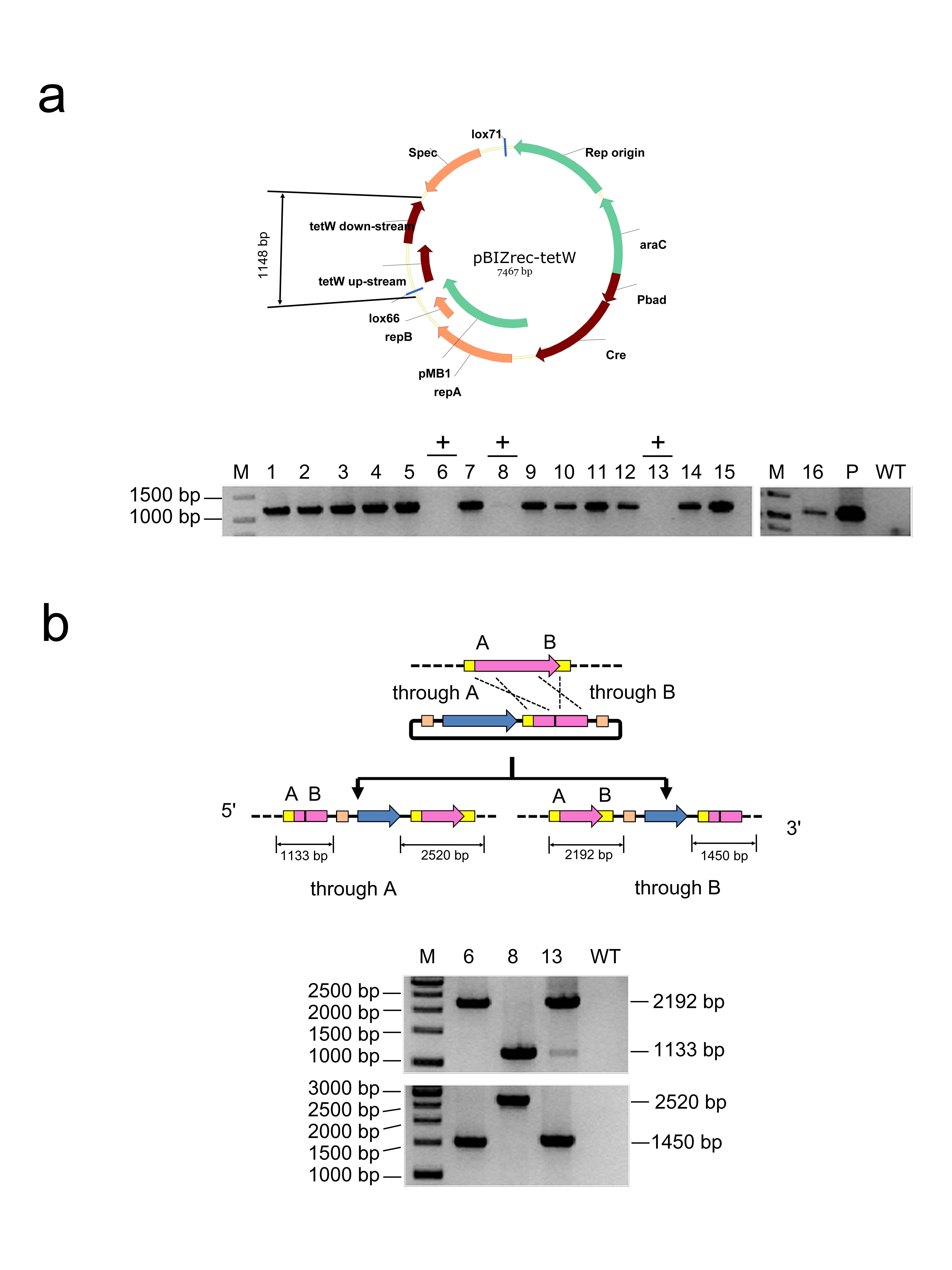

Supplementary Figure 4. Deletion of *tetW* gene in *B. longum* IF3-53 by IPSD assisted genome engineering. (a) Single crossover integration of *tetW-Smr* fragment into the chromosome of *B. longum* IF3-53. Colony PCR using primers pBrec-F and pBrec-R was performed to detect the intact plasmid pBIZrec-tetW. Three colonies without the corresponding band were regarded as candidate single crossover clones, which were labelled "+". (b) The three candidate recombinants were further confirmed by DNA extraction and PCR using primers tetWleft-F and pBrecSC-R (upper gel), or pBrecSC-F and tetWright-R (lower gel). M, GeneRuler^TM^ 1kb DNA ladder; P, DNA of *B. longum* IF3-53 harboring plasmid pBIZrec-tetW; WT, DNA of wild type *B. longum* IF3-53.

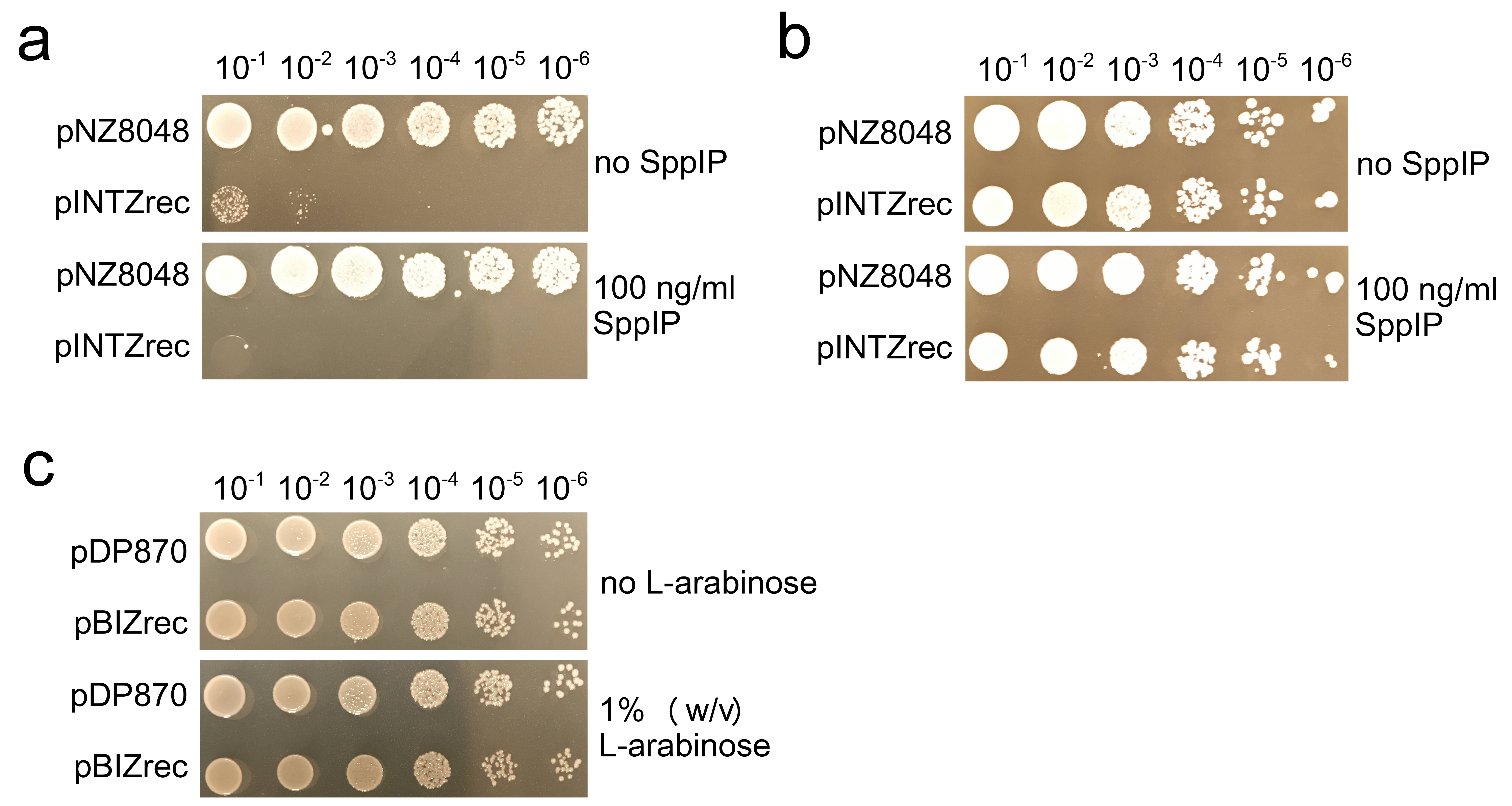

Supplementary Figure 5. The inducible promoters used to drive the expression of recombinase are not effective in some strains. Growth of the indicated *L. sakei* NC03 strain (a) and *L. rhamnosus* GG strain (b) on MRS agar plate supplemented with 10 μg/ml chloramphenicol and in the presence or absence of 100 ng/ml SppIP. (c) Growth of the indicated *B. lactis* Bb12 strain on MRSc agar plate supplemented with 100 μg/ml spectinomycin and in the presence or absence of 1 % (w/v) L-arabinose. The results show that in *L. sakei* NC03, plasmid pINTZrec was destructed even without of additional inducer. While in *L. rhamnosus* GG and *B. lactis* Bb12, the inducers could not induce plasmid self-destruction.

Supplementary Table 1. Strains and plasmids used in this study.

| Strain or plasmid | Characteristics ^a^ | Source |
| --- | --- | --- |
| Strains |  |  |
| *E. coli* DH5α | *fhuA2 Δ(argF-lacZ)U169 phoA glnV44 Φ80 Δ(lacZ)M15 gyrA96 recA1 relA1*  *endA1 thi-1 hsdR17* | Invitrogen |
| *E. coli* VE7108 | Km^r^, host of pNZ8048 | ^1^ |
| *L. gasseri* DSM 14869 | Human vaginal isolate | ^2^ |
| *L. paracasei* BL23 | Origin unclear | ^3^ |
| *L. plantarum* NL42 | Cheese isolate | ^4^ |
| *L. acidophilus* ATCC 4356 | Human pharynx isolate | ^5^ |
| *L. sakei* NC01 | Sausage isolate | This study |
| *L. sakei* NC03 | Sausage isolate | This study |
| *B. longum* NCC 2705 | Human feces isolate | Nestlé Culture Collection |
| *B. longum* IF3-53 | Infant feces isolate | ^6^ |
| *B. lactis* Bb12 | Commercial probiotic | Chr. Hansen Ltd. (Hørsholm, Denmark) |
| *L. gasseri* DSM 14869 *Δupp* | *L. gasseri* DSM 14869 *upp* deletion mutant | This study |
| *B. longum* IF3-53 *ΔtetW* | *B. longum* IF3-53 *tetW* deletion mutant | This study |
| *B. longum* NCC 2705 P*_hup_::LpKatL* | *B. longum* NCC2705 containing integrated *LpKatL* gene under the control of P_hup_ promoter | This study |
| Plasmids |  |  |
| pNZ8048 | Cm^r^, SH71 replicon plasmid | ^7^ |
| pDP870 | Sm^r^, *E. coli-B. longum* shuttle cloning vector | ^8^ |
| pEM76 | Source of six sequence | ^9^ |
| pEM94 | Source of *β*-recombinase gene | ^10^ |
| pVPL3017 | Cm^r^, pSIP411 derivative, containing sakacin-inducible expression element | ^11^ |
| pAdTrack-Cre | Source of *cre* gene | ^12^ |
| pDP401-LpKatL | pDP870 derivative, containing *LpKatL* gene | ^13^ |
| pVPL3017-rec | pVPL3017 derivative, with *recT1* replaced with *β*-recombinase gene | This study |
| pNZ8048-SIX | pNZ8048 derivative containing two six DNA fragments flanking Cm^r^ gene | This study |
| pNZmcs-SIX | pNZ8048-SIX derivative containing a multiple-cloning site | This study |
| pINTZrec | pNZmcs-SIX derivative containing *β* recombinase gene expression cassette | This study |
| pINTZrec-Δupp | pITZrec derivative containing upstream and downstream DNA of *L. gasseri* DSM 14869 *upp* gene | This study |
| pDP870-lox | pDP870 derivative containing lox69 and lox71 sequences flanking Sm^r^ gene | This study |
| pBIZrec | pDP870-lox derivative containing *cre* recombinase gene expression cassette | This study |
| pBIZrec-∆tetW | pBIZrec derivative containing upstream and downstream DNA of *B. longum* IF3-53 *tetW* gene | This study |
| pBIZrec-P_hup_-Kat | pBIZrec derivative containing *LpKatL* gene placed under the control of P*_hup_* from *B. longum* NCC 2705 by a polycistronic operon structure | This study |

^a^ Km^r^, kanamycin resistance; Cm^r^, chloramphenicol resistance; Sm^r^, spectinomycin resistance.

Supplementary Table 2. Oligonucleotides used in this study.

| Oligonucleotides | Sequence, 5'-3'^a^ | Restriction sites |
| --- | --- | --- |
| SIX-F1 | GACCGGTCGACAATTATTAGGGGGAGAAG | *Sal*I |
| SIX-R1 | CGCCAGTCGACGAGTCGTGCATAACCAAT | *Sal*I |
| SIX-F2 | GACCGCTGCAGAATTATTAGGGGGAGAAG | *Pst*I |
| SIX-R2 | CGCCAAAGCTTGAGTCGTGCATAACCAAT | *Hind*III |
| linker-F | GATCTGAGCTCATGCATGGGCCCGATCGCTAGCGGCCGCATGCGGATCCTGCA |  |
| linker-R | GGATCCGCATGCGGCCGCTAGCGATCGGGCCCATGCATGAGCTCA |  |
| upp-up-F | TAATTGGGCCCAAATAATGGAAACTAAGATT | *Apa*I |
| upp-up-R | GTTCAGCATGCTAACAAGAGCTCAGATAAATGTTTCTTAAATCGT | *Sph*I, *Sac*I |
| upp-down-F | CAGGAGAGCTCTTGTTCGGATCCAAGTAATTTTACTCAAAAATCT | *Sac*I, *BamH*I |
| upp-down-R | TTACAGCATGCAAAACGCAAATTACAGGAAGAG | *Sph*I |
| uppleft-F | TTACCAGATTTTGAAATTGAGTT |  |
| uppright-R | AGTAAAGCGTATCTCCTAACTCT |  |
| pIrec-F | AGATTTATTGAGAGGAGGGATTATT |  |
| pIrec-R | CGTTTGTTGAACTAATGGGTGCT |  |
| pIrecSC-F | AAAGTTTTCGGGCTACTCTCTCCT |  |
| pIrecSC-R | GGAATTGTCAGATAGGCCTAATGACT |  |
| uppseq-F | GAACAATTAGTCCTGCTTATATG |  |
| uppseq-R | CGACTACAGATTTCTCATTCACT |  |
| Lox66-F | ATTTGATATCTACCGTTCGTATAATGTATGCTATACGAAGTTATCTCGAGCTCATATGCATGGGC | *EcoR*V |
| Lox71-R | CGGGGAATTCTACCGTTCGTATAGCATACATTATACGAAGTTATCTGGCGAATGGCGATTTTCGT | *EcoR*I |
| araP-Cre-F1 | ACTTGGTCTACGCTTGCATAATGTGCCTGTCAAAT | *Acc*I |
| araP-Cre-R1 | CGGTCAGTAAATTGGATACCATGGTTAATTCCTCCTGTTAGCCC |  |
| araP-Cre-F2 | GGGCTAACAGGAGGAATTAACCATGGTATCCAATTTACTGACCG |  |
| araP-Cre-R2 | ATGCAAGCTTCTAATCGCCATCTTCCAGCAG | *Hind*III |
| tetW-up-F | TGTAGGGGCCCAGTTTTTTGTACCCAATTTAAG | *Apa*I |
| tetW-up-R | TCTTCTGAAACATATAGCGCACCT |  |
| tetW-down-F | TGATCTCCGCTAAAGATGGCCACCACTCCCGCTTGGTTCCGGTGT |  |
| tetW-down-R | GACTGGCTAGCCGACAGCGGCCTGATACCCTT | *Nhe*I |
| tetWleft-F | CTTATAGTGGGAACAAAGGATTATGATAG |  |
| tetWright-R | AACATATGGCGCACCTTGTCCAG |  |
| pBrecSC-F | CACTCTCAACTCCTGATCCAAACAT |  |
| pBrecSC-R | ATTTGGGAAATATTCATTCT |  |
| hup-up-F | CCTGCGGGCCCTGCGCAAATCCGGCTATCG | *Apa*I |
| hup-up-R | TCCATAAGGACTCCTTAAGGTCACTCGGTGACGGCCT |  |
| hup-down-F | ATATTATCAGTGATTAACTGCTCGTAGCGATTACTTCG |  |
| hup-down-R | TCATCGCTAGCCAGCTGTACACCGGTGCCG | *Nhe*I |
| Kat-F | AGTGACCTTAAGGAGTCCTTATGGAAATGACGGAAAAATT |  |
| Kat-R | CGCTACGAGCAGTTAATCACTGATAATATCAGCAAT |  |
| Katleft-F | CAGCCGGCGTGGGCCCCTCGTT |  |
| Katright-R | CTGGTCCGCCGCTTCGTGCCGTT |  |
| pBrec-F | CACTCTCAACTCCTGATCCAAACAT |  |
| pBrec-R | AGTGAGCGCAACGCAATTAATGT |  |

^a^ The restriction sites are underlined.
